# A field-potential signature of pre-burst dynamics in human higher-order thalamic neurons

**DOI:** 10.64898/2026.11.13.766342

**Authors:** L Varisa, E Nester, B Bhattarai, A Sridharan, KP Rains, Z Wrobel, J Hong, R Suri, LL Lehnhoff, DP Harris, S Vladimir, K Choi, CM Schusse, Z Mirzadeh, ST Foldes, AI Yang

## Abstract

The aperiodic (1/f) component of the local field potential (LFP), long dismissed as noise, is increasingly being recognized as physiologically meaningful. Its exponent has been proposed to index excitation:inhibition (E:I) balance, with higher exponents (steeper spectra) corresponding to greater inhibition. Yet this hypothesis remains debated and has seldom been tested against single-neuron activity. Here, we used rare concurrent single-neuron and LFP recordings from the human pulvinar during goal-directed behavior and quiet wakefulness. Pulvinar recordings in both states revealed signatures of low-threshold calcium spikes (LTS) bursting. Prior work with intracellular recordings has established that LTS bursts require a sustained, spike-free period of hyperpolarization followed by a triggering depolarization, offering a physiologically-grounded test of the E:I hypothesis. We found that the exponent was higher during pre-burst silence relative to duration-matched silent windows preceding tonic spikes. The exponent also reflected the graded nature of burst-priming hyperpolarization: bursts with more constituent spikes followed longer silences and higher exponent peaks. Whereas these relationships held in both behavioral states, co-bursting across neurons exceeded chance only during goal-directed behavior. Co-bursts occurred on a slow (∼100 ms) timescale, consistent with a shared cortical input mechanism, and were preceded by a larger rise in the exponent. Together, by linking a field potential measure to a known cellular mechanism, our findings show that the aperiodic exponent tracks the graded priming of thalamic bursts, providing human single-neuron support for the E:I hypothesis.

## INTRODUCTION

Historically, research on brain electrical activity has focused on oscillatory components. However, beneath these oscillations lies an aperiodic component of the local field potential (LFP), which is used as a window into brain state (Gao, Peterson and Voytek, 2017; Donoghue *et al*., 2020). LFP power decreases with frequency, and the absolute value of the slope of this decline is termed the aperiodic exponent (Miller *et al*., 2009; Donoghue *et al*., 2020). The exponent is believed to steepen as arousal decreases, such as during sleep or anesthesia (Lendner *et al*., 2020).

Steeper spectra have been interpreted as reflecting relatively greater inhibition, and flatter spectra relatively greater excitation, which has led to the exponent being discussed as a possible index of E:I balance (Gao, Peterson and Voytek, 2017; Donoghue *et al*., 2020). However, most supporting evidence comes from computational models or comparisons between brain states (Gao, Peterson and Voytek, 2017; Lendner *et al*., 2020).

Previous studies of the neuronal correlates of aperiodic slope are limited to subthalamic recordings from rodent models of Parkinson’s disease, which showed that periods of low firing coincide with a higher exponent (Wiest *et al*., 2023). Thalamic cells fire in two modes: tonic spiking or bursts (Sherman, 2001; Sherman and Guillery, 2002). In intracellular recordings, firing mode follows membrane potential: depolarization inactivates T-type calcium channels and yields tonic spiking, whereas sustained hyperpolarization de-inactivates them, so that a subsequent depolarizing input triggers low-threshold spikes (LTS) bursting (Jahnsen and Llinás, 1984; Sherman, 2001; Sherman and Guillery, 2002). Bursting is more commonly observed during sleep than in wakefulness, but it also occurs in awake animals (Ramcharan, Gnadt and Sherman, 2000) and is more prevalent in higher-order than in first-order thalamic nuclei(Ramcharan, Gnadt and Sherman, 2005).

We studied concurrent single-unit and LFP recordings from the human pulvinar during both quiet wakefulness and goal-directed behavior. As thalamic bursts represent hyperpolarization-primed neuronal activity, we hypothesized that the exponent should be higher during the silent period preceding a burst.

## RESULTS

We recorded single-units from the pulvinar in the awake rest state (4 subjects, 8 recordings, 95 isolated single-units) and while performing a cognitive task (Fig. 1a,b; 10 subjects, 119 isolated single-units). Bursts were detected using published criteria (Lu, Guido and Sherman, 1992; Ramcharan, Gnadt and Sherman, 2005) and analyses were restricted to single-units with at least 10 burst events: 28.4% (27/95) of rest and 38.7% (46/119) of task neurons. Consistent with prior literature (Boshra *et al*., 2026), burst rates were higher at rest (6.74 ± 2.03 bursts/min; mean ± s.e.m., n = 27 neurons) than during task performance (2.24 ± 0.45 bursts/min; n = 46 neurons; two-sided Wilcoxon rank-sum test, p = 0.0009).

**Figure 1:**
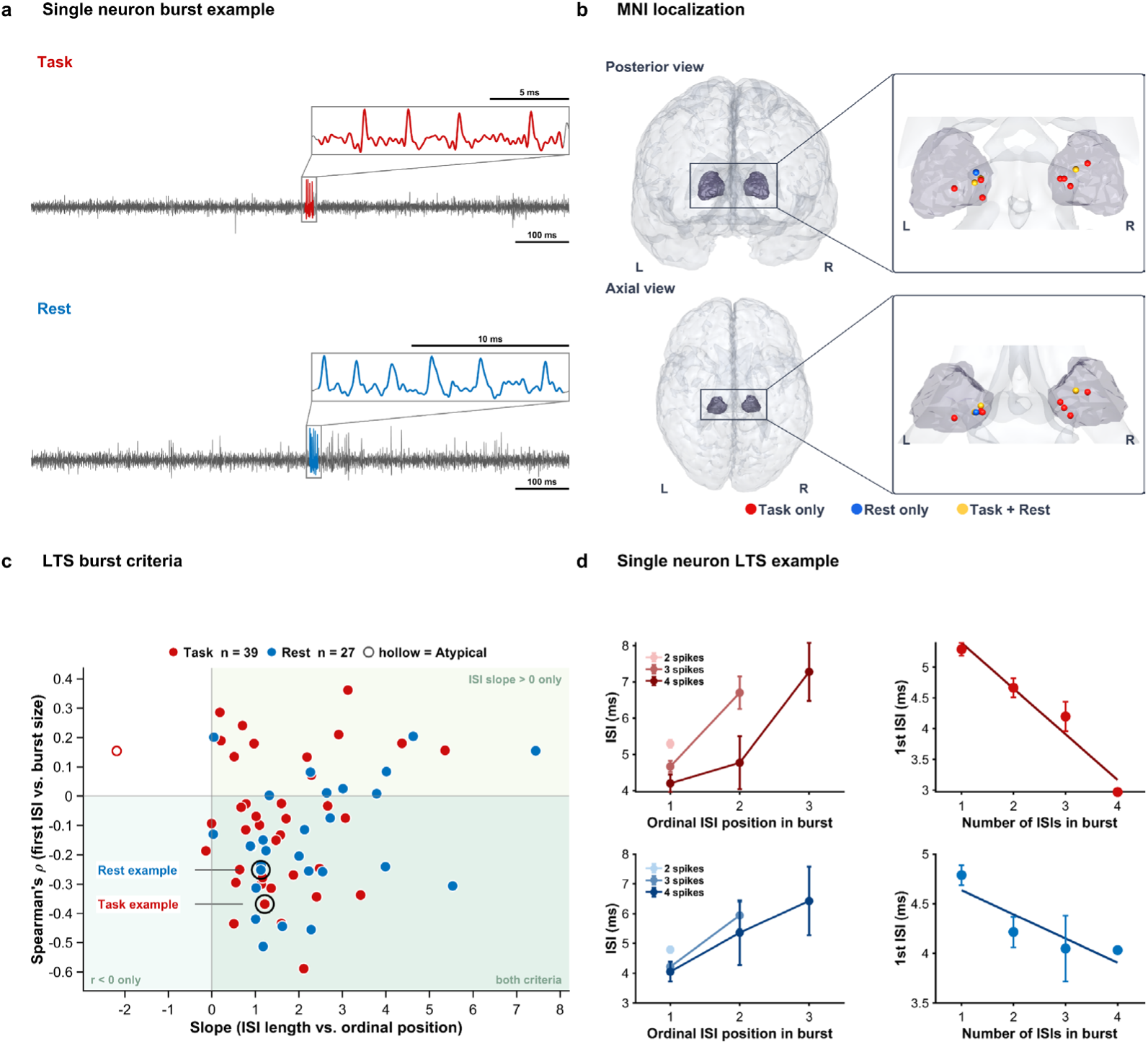
Putative LTS bursting neurons from the human pulvinar in two states. a. Example single-unit bursts from a task neuron (red) and rest neuron (blue). The inset magnifies the burst. b. MNI localization of pulvinar units for Task (red), Rest (blue), and Task + Rest (yellow) subjects c. LTS burst criteria for each neuron. The x-axis shows the slope of the inter-spike interval (ISI) across ordinal position within bursts (ms/position). The y-axis shows the per-neuron correlation between the first ISI and burst size. Shaded quadrants mark the criteria: positive ISI slope only (upper right), r < 0 only (lower left), and both criteria (lower right). Filled spheres meet both criteria, and hollow spheres mark atypical neurons. Circled points mark the example task and rest neurons shown in (d). d. Example single-neuron bursts from a task neuron (top, red) and a rest neuron (bottom, blue). Left: ISI (ms) as a function of ordinal ISI position within the burst, plotted separately for bursts of 2, 3, and 4 spikes. ISIs lengthen across the burst, as expected for low-threshold-spike bursts. Right: Duration of the first ISI (ms) as a function of the number of ISIs in the burst. Larger bursts start with a shorter first ISI.

To test the E:I hypothesis, we first confirmed that detected bursts showed features of low-threshold spike (LTS) bursts. Within bursts we tested two criteria: (i) ISIs lengthen with ordinal position, and (ii) bursts with a longer first ISI contain fewer spikes (Domich, Oakson and Steriade, 1986; Ramcharan *et al*., 2000) (Fig. 1c,d). For each neuron, we quantified (i) as the slope of ISI length against ordinal position and (ii) as the Spearman correlation (ρ) between first-ISI length and burst size (Fig. 1c). Single-neuron examples from task (red) and rest (blue) show both features: ISIs increased with ordinal position (Fig. 1d, left), and bursts with longer first ISIs contained fewer spikes (Fig. 1d, right). All bursting neurons in the task recordings (46/46) and 26 of 27 (96.3%) in the rest recordings met at least one criterion.

### Aperiodic exponent reflects increased inhibition during pre-burst silence leading to low-threshold burst firing

LTS bursting occurs when the membrane is hyperpolarized, which de-inactivates T-type Ca²⁺ channels, whereas neurons fire tonically when more depolarized (Jahnsen and Llinás, 1984; Sherman, 2001; Sherman and Guillery, 2002). The aperiodic exponent of the LFP is a proxy for E:I balance, where steeper slopes indicate more inhibition (Gao, Peterson and Voytek, 2017; Donoghue *et al*., 2020). From that, we predicted that the silent period before a burst should be steeper than the silent period before tonic firing. Intracellular recordings have shown that LTS bursts follow a prolonged period (50-100 ms) of hyperpolarization leading to bursting (Jahnsen and Llinás, 1984; Gutierrez *et al*., 2001). As a putative metric of E:I balance, a high aperiodic exponent is thought to index increased inhibition, whereas a low aperiodic exponent is thought to index increased excitation. Note that the aperiodic component can be linearly fitted in log-log space with a negative slope. Correspondingly, a steeper (flatter) slope is thought to index greater inhibition (excitation) (Lendner *et al*., 2020). An example average PSD fit across the full spectrum and the 10–50 Hz fit range is shown for the pre-burst, burst-peak and tonic windows (Fig. 2b, inset).

**Figure 2:**
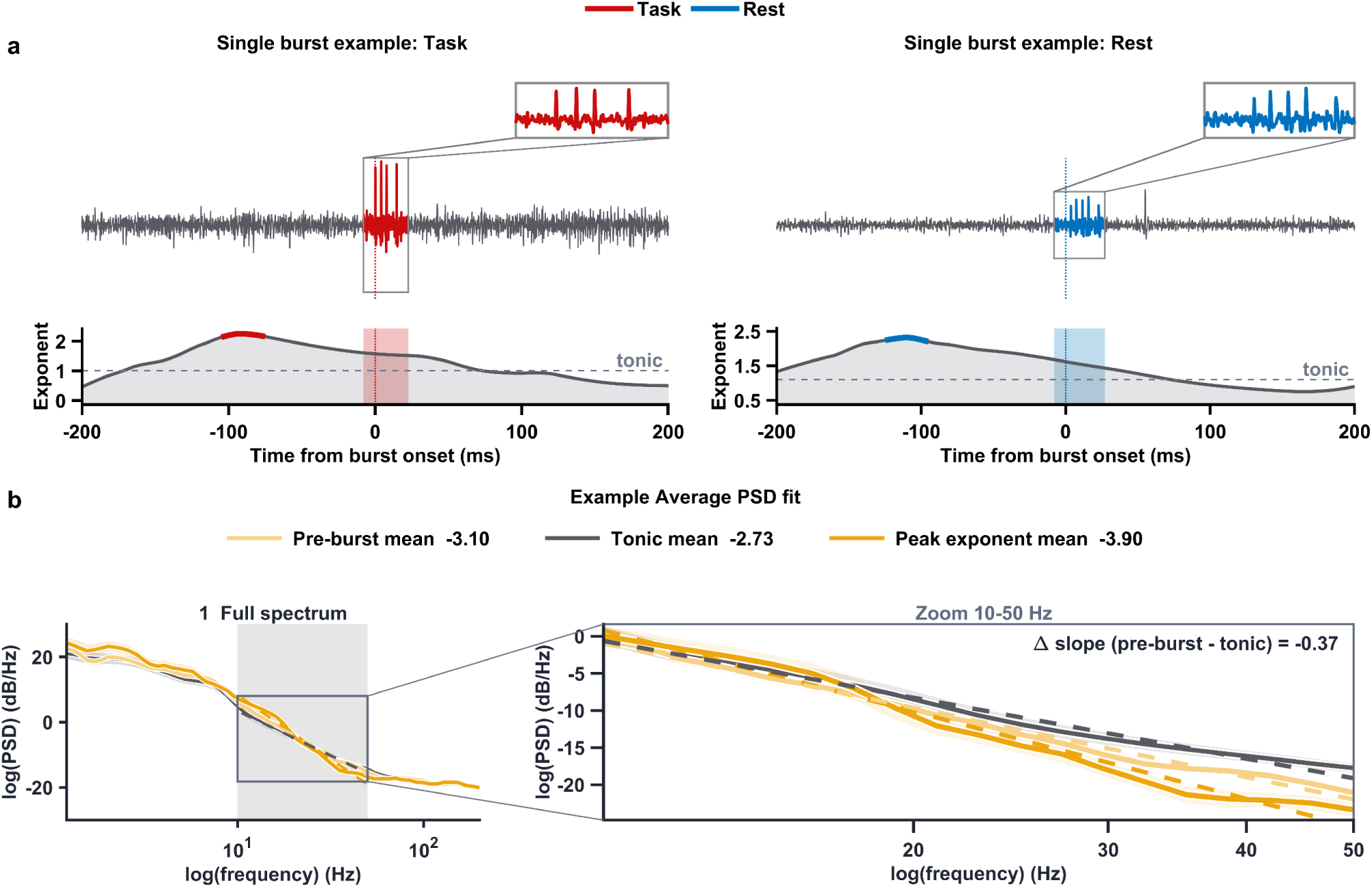
Pulvinar LTS bursting as a testbed for the E:I hypothesis of aperiodic exponent. a. Single burst examples. Raw trace (top) and aperiodic exponent (bottom) around one burst, aligned to burst onset, for an example task neuron (red) and rest neuron (blue). Below each trace, the time-resolved aperiodic exponent is plotted against time from burst onset (ms), with the tonic baseline shown as a dashed line. The shaded box marks the burst window. b. Average PSD fit example for a single neuron. The curves show the average tonic (grey), average pre-burst (light), and average peak exponent (full color). Zoom on 10–50 Hz showing the spectral slopes (solid lines, data; dashed, fit)

In single-neuron examples, the exponent increased during the pre-burst silent period and returned to its tonic level afterward (Fig. 2a).The silent period was defined as an interval in which the neuron fired no spikes for at least 100 ms before burst onset, consistent with the hyperpolarization duration required to de-inactivate T-type Ca²⁺ channels (Lu, Guido and Sherman, 1992; Ramcharan, Gnadt and Sherman, 2005). To test this effect across neurons, we normalized time bins in the silent period to account for variable pre-burst durations such that−100% marks the onset of the silent period (the last spike preceding the silence) and 0% marks burst onset (the first spike of the burst). We then compared pre-burst windows with randomly sampled pre-tonic windows matched for silent-period duration using a cluster-based permutation test. (Fig. 3a; labels shuffled within neurons, 1,000 permutations, two-sided p<0.05)(Maris and Oostenveld, 2007). The aperiodic exponent was higher before bursts during both task performance ([−78%,−1%] of the silent period; cluster p = 0.021) as well as in the rest state ([−100%, −0%]; cluster p = 0.0049).

**Figure 3:**
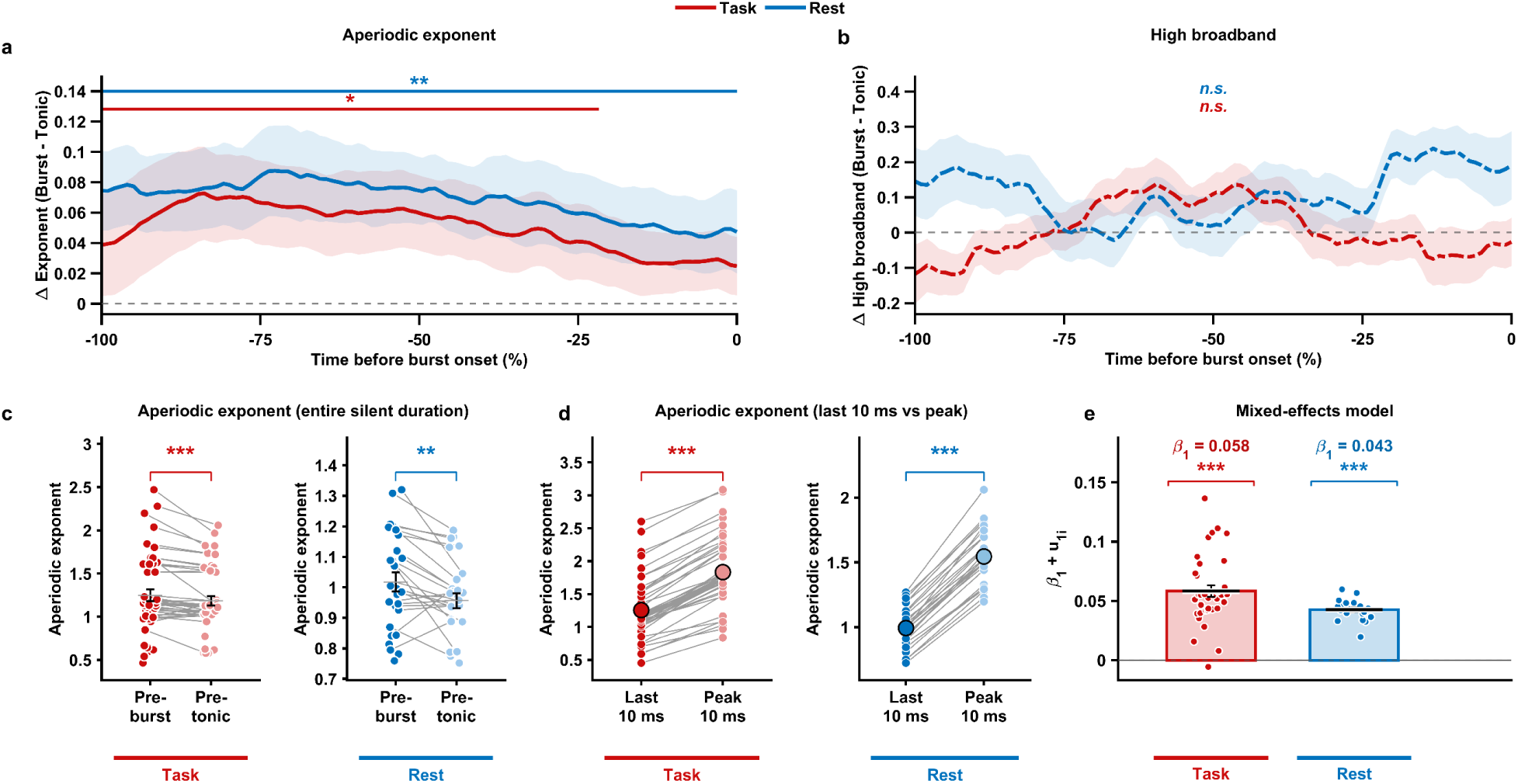
Aperiodic exponent reflects hyperpolarization-induced bursting. a. Pre-burst vs. pretonic aperiodic exponent: change in exponent (preburst - pretonic) as a function of time before burst onset (-100% to 0%) for task(red) and rest(blue). Horizontal bars mark time windows that differ significantly from zero (cluster permutation test; Task *, Rest **). b. Pre-burst vs. pretonic high broadband power: change within high-frequency broadband power, which was not significant in either state (n.s.). c. Mean aperiodic exponent across the entire silent duration, paired by unit d. Average aperiodic exponent within the last 10ms of the pre-burst vs. 10ms of peak value, paired by unit e. Mixed-effects model of the exponent slope over the pre-burst period. Bars show the fixed-effect slope (Task β₁ = 0.056, n = 36; Rest β₁ = 0.043, n = 22; both ***) and dots show unit-level slopes (β₁ + uᵢ). *p < 0.05, **p < 0.01, ***p < 0.001; n.s., not significant.

If the rise in aperiodic exponent represents hyperpolarization, then the depolarization triggering the burst should manifest in decrease of the aperiodic slope immediately before the burst. In fact, we found that aperiodic exponent in the last 10ms before the spike onset was lower than the peak aperiodic exponent during the preceding silence (task: 1.256 ± 0.067 vs. 1.833 ± 0.075, mean ± s.e.m., n = 46 neurons; two-sided Wilcoxon signed-rank test, p = 3.5 × 10⁻⁹; rest: 0.994 ± 0.030 vs. 1.545 ± 0.041, n = 27 neurons; p = 5.6 × 10⁻⁶; Fig. 3c). The decrease was present in all 46 task neurons and all 27 rest neurons. Neuron-wise, time-averaged aperiodic exponent was greater during the pre-burst silence than during matched pre-tonic windows (paired, within-neuron comparison; one-sided Wilcoxon signed-rank test). During task performance, the exponent was 1.248 ± 0.067 (mean ± s.e.m.) before bursts and 1.184 ± 0.055 before tonic spikes (n = 46 neurons; exponent higher before bursts in 35 of 46 neurons; p = 3.4 × 10⁻⁴). At rest, it was 1.018 ± 0.031 before bursts and 0.956 ± 0.025 before tonic spikes (n = 27 neurons; higher during pre-bursts in 20 of 27 neurons; p = 1.0 × 10⁻³; Fig. 3d).

To demonstrate the specificity of these findings, a parallel analysis was performed with High-frequency broadband power (70-200Hz) (Saez et al., 2018), a marker of local population firing (Manning *et al*., 2009). Pre-Burst and tonic periods did not differ within task or rest recordings for high broadband (two-sided Wilcoxon signed-rank test, P = …; Fig. 3b). That is, differential pre-burst aperiodic exponent dynamics across firing modes were distinguishable dynamics from high-frequency broadband power (offset of power spectra).

Although the LFP primarily reflects postsynaptic currents (Buzsáki, Anastassiou and Koch, 2012), it is also influenced by neuronal firing, including rate and synchrony (Manning *et al*., 2009), and the aperiodic component itself has been linked to population spiking (Manning *et al*., 2009). As pre-burst and tonic silence periods were defined with respect to the presence of firing within each neuron, we performed further analysis to assess whether our findings were robust to differences in single unit activity of the remaining simultaneously recorded neurons. To do so, we fitted a linear within-between mixed-effects model (Bates *et al*., 2015) to predict aperiodic exponent across pre-burst and pre-tonic windows. Burst vs. tonic remained a significant predictor of the aperiodic exponent after controlling for firing rate and pairwise synchrony of the remaining neurons, with a positive effect in both states (task: β₁ = 0.058, 95% CI [0.028, 0.089], p = 1.6 × 10⁻⁴; rest: β₁ = 0.043, 95% CI [0.003, 0.082], p = 0.035). Per-neuron effects (fixed effect + random effect of pre-burst vs. tonic, β₁ + u₁ᵢ) were positive in both states (task: 0.058 ± 0.005, n = 36 neurons; rest: 0.043 ± 0.002, n = 22 neurons; mean ± s.e.m.; Fig. 3e), indicating that the effect was consistently observed across neurons.

### Strength of pre-burst Aperiodic Exponent increase correlates with strength of burst

Variability in the strength of pre-burst hyperpolarization has been associated with the strength of individual burst events as the recovery of T-type channels from inactivation grows with hyperpolarization depth and duration (Jahnsen and Llinás, 1984; Alitto *et al*., 2019). We tested this in neurons with bursts of varying size. For each burst, we measured the preceding silence duration, the peak exponent during that silence, and the burst size (spikes per burst) (Fig. 4a), and computed Spearman correlations across burst events within neurons(Benjamini and Hochberg, 1995).

**Figure 4:**
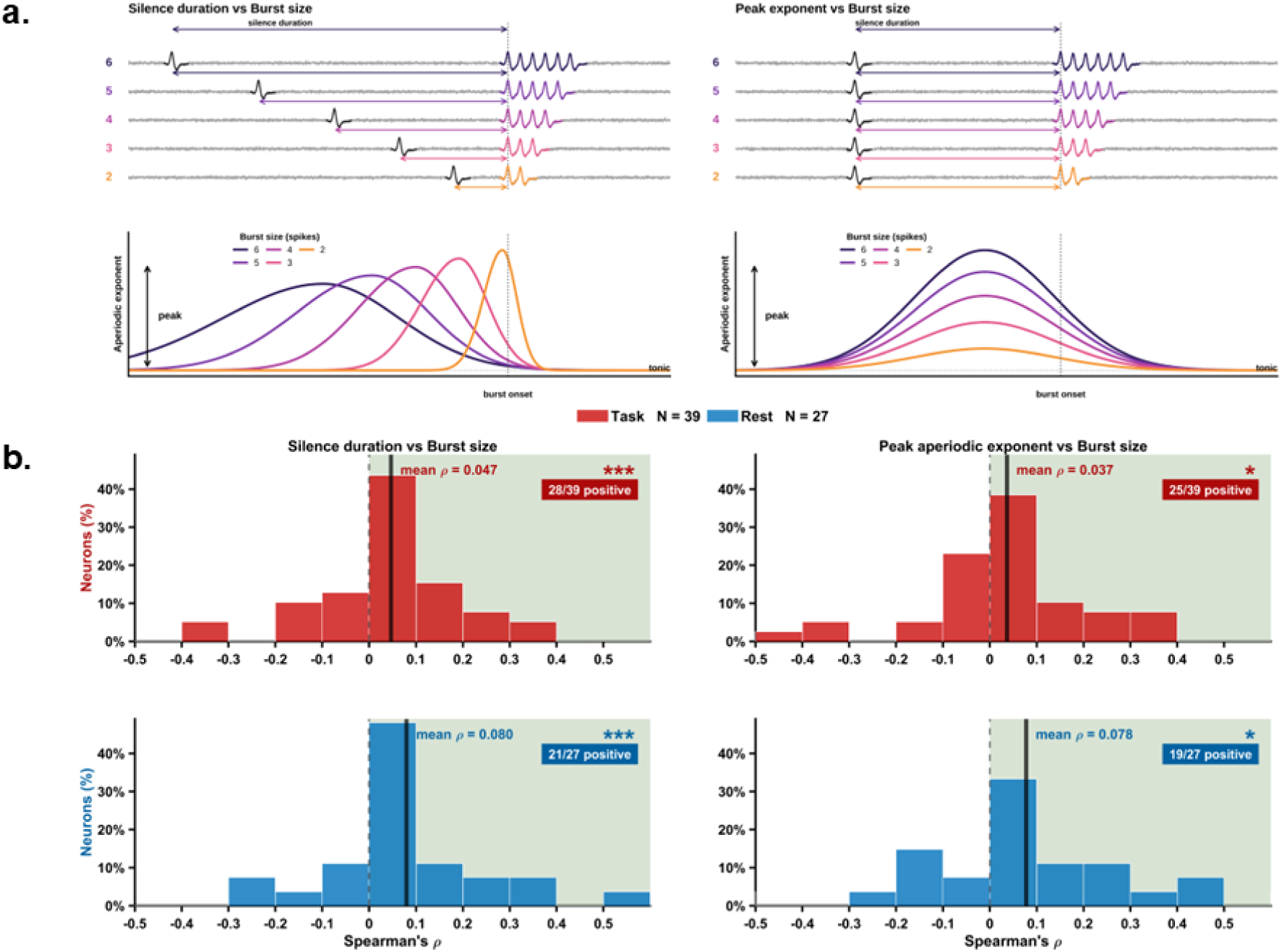
Aperiodic exponent reflects graded bursting. a. Left: Schematic of bursts of 2–6 spikes, each preceded by a silent period. Right: Schematic of predicted aperiodic exponent (AE) before burst onset. Larger bursts (darker purple) follow longer silent periods and show a higher peak exponent relative to the tonic baseline. b. Correlations: Left: Distribution across neurons of the Spearman correlation (r) between silent-period duration and burst spike count. Right: Distribution of the Spearman correlation between peak exponent and spike count. Top, Task (red, N = 39); bottom, Rest (blue, N = 27). Solid vertical lines mark the mean ρ, dashed lines mark ρ = 0, and the shaded region marks positive correlations. The text gives the mean ρ and the number of neurons with positive ρ. *p < 0.05, ***p < 0.001.

Longer silences predicted larger bursts in 28 of 39 task neurons (71.8% vs. 28.2% with non-positive correlations; mean ρ = 0.047 ± 0.024 s.e.m.; Fisher’s exact test, FDR-corrected p = 4.8 × 10⁻⁴) and in 21 of 27 rest neurons (77.8% vs. 22.2%; mean ρ = 0.080 ± 0.031s.e.m.; Fisher’s exact test, FDR-corrected p = 1.9 × 10⁻⁴; Fig. 4b). P-values were corrected using the Benjamini– Hochberg procedure (α = 0.05) across the two predictors (silence duration and peak aperiodic exponent) within each condition.

The exponent therefore not only is distinguishable between pre-burst and tonic periods, but reflects the graded effect between the strength of hyperpolarization and how strongly a neuron bursts.

### Slow timescale co-bursting during task behavior

As thalamic nuclei, including pulvinar, lack local recurrent excitatory connectivity unlike in cortex, co-bursting emerges from shared common input from cortical regions (Liu *et al*., 2026). Example co-bursting events are shown in Fig. 5a. We tested whether co-bursts occur more frequently than expected by chance analyzing neurons recorded simultaneously with the same microwire bundle (Fig. 5b).

**Figure 5:**
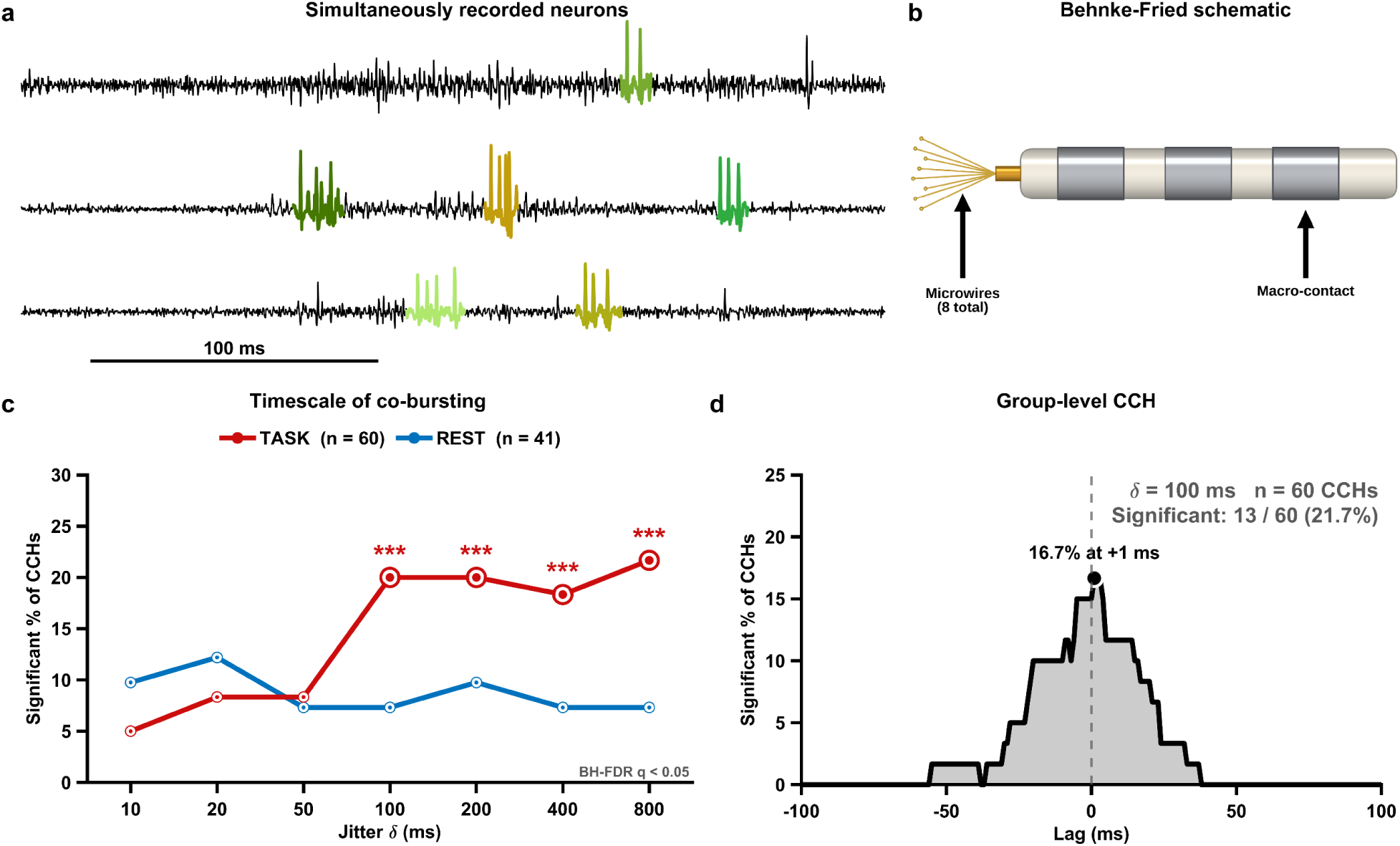
Task performance recruits co-bursting with timescales consistent with shared cortical input. a. Example traces of simultaneously recorded neurons from the same wire bundle b. A schematic of the Behnke-Fried electrode showing the microwires and macro-contacts. c. Timescale of co-bursting. Percentage of neuron pairs with significant co-bursting as a function of δ (ms) for Task (red, n = 60) and Rest (blue, n = 41). Task neurons show elevated co-bursting at jitter windows of 100 ms and above (***, FDR-corrected). d. Group-level CCH (cross-correlogram) of burst times across pairs, with the peak at a lag of +/-50 ms showing the percentage of CCHs with a significant cluster at each lag.

For each neuron, we built a corresponding population train from the burst onsets of all other simultaneously recorded neurons. We then constructed a conditional discharge-rate cross-correlogram (CCH, lag [-2, 2]s) between each single unit-level burst train and the corresponding population burst train (60 and 41 CCHs during task performance and awake rest state, respectively; see Methods).

We assessed statistical significance of CCH peaks using a jitter-based permutation test(Liu *et al*., 2026), whereby the population level burst train was circularly shifted in time by a random offset drawn uniformly from −δ to +δ, generating 1,000 surrogate per CCHs per neuron pair. A CCH was considered to have a significant peak if the observed count within ± 2 s of zero lag exceeded the 97.5th percentile of the surrogate distribution. Shifts of up to ±δ disrupt coordination at timescales finer than δ while preserving slower structure, so repeating the test across δ = 10, 20, 50, 100, 200, 400 and 800 ms reveals the timescale of co-bursting: a significant peak at a given δ indicates excess co-bursting at timescales finer than δ.

During task performance, the percentage of statistically significant CCHs was greater than the theoretical chance level (5%; FDR-corrected threshold across jitter values: 13.3%, i.e., ≥ 8 of 60 CCHs) from δ = 100 to 800 ms (11–13 of 60 CCHs, 18.3–21.7%; q = 9.6 × 10⁻⁵ to 6.0 × 10⁻⁴; Fig. 5c), but not at δ ≤ 50 ms (3–5 of 60, 5.0–8.3%; q ≥ 0.28). In the awake rest state, the percentage of significant CCHs did not exceed the corrected threshold (17.1%, i.e., ≥ 7 of 41 CCHs) at any δ (3–5 of 41, 7.3–12.2%; q ≥ 0.15), indicating that co-bursting was detectable during goal-directed behavior but not at rest. Detection of co-bursting on a timescale of ∼100 ms indicates that it occurs on a timescale of tens of milliseconds, slower than millisecond-precise synchrony, consistent with shared input as an underlying mechanism. At the group level, statistically significant CCH peaks showed co-bursting within −56 to +38 ms (Fig. 5d; δ = 100 ms). We therefore identified individual co-burst events as temporal epochs in which at least two simultaneously recorded neurons burst within 50 ms of each other.

The aperiodic exponent was higher during the pre-co-burst silence than during matched pre-tonic windows ([−84%, −34%] of the silent period; p = 0.012; Fig. 6a). Here, silent windows were defined as a period without spikes in any of the concurrently-recorded neurons within each population. Similarly, broadband power was not different between pre-co-burst and tonic windows (Fig. 6b). As LFP is a population-level signal, we expected that the magnitude of aperiodic exponent increase during the pre-co-burst windows would be greater vs. the magnitude of aperiodic exponent increase during silent periods determined at the single neuron level. In fact, the average aperiodic exponent during pre-co-burst windows was greater - consistent with greater hyperpolarization - compared to the single-neuron pre-burst windows 0.31 ± 0.17 vs. 0.07 ± 0.02, mean ± s.e.m.; one-sided Wilcoxon rank-sum test, W = 267, p = 0.021; Fig. 6c). We confirmed that pre-tonic periods used in each analysis were not different (1.12 ± 0.22 vs. 1.18 ± 0.06; two-sided Wilcoxon rank-sum test, W = 165, p = 0.537).

**Figure 6:**
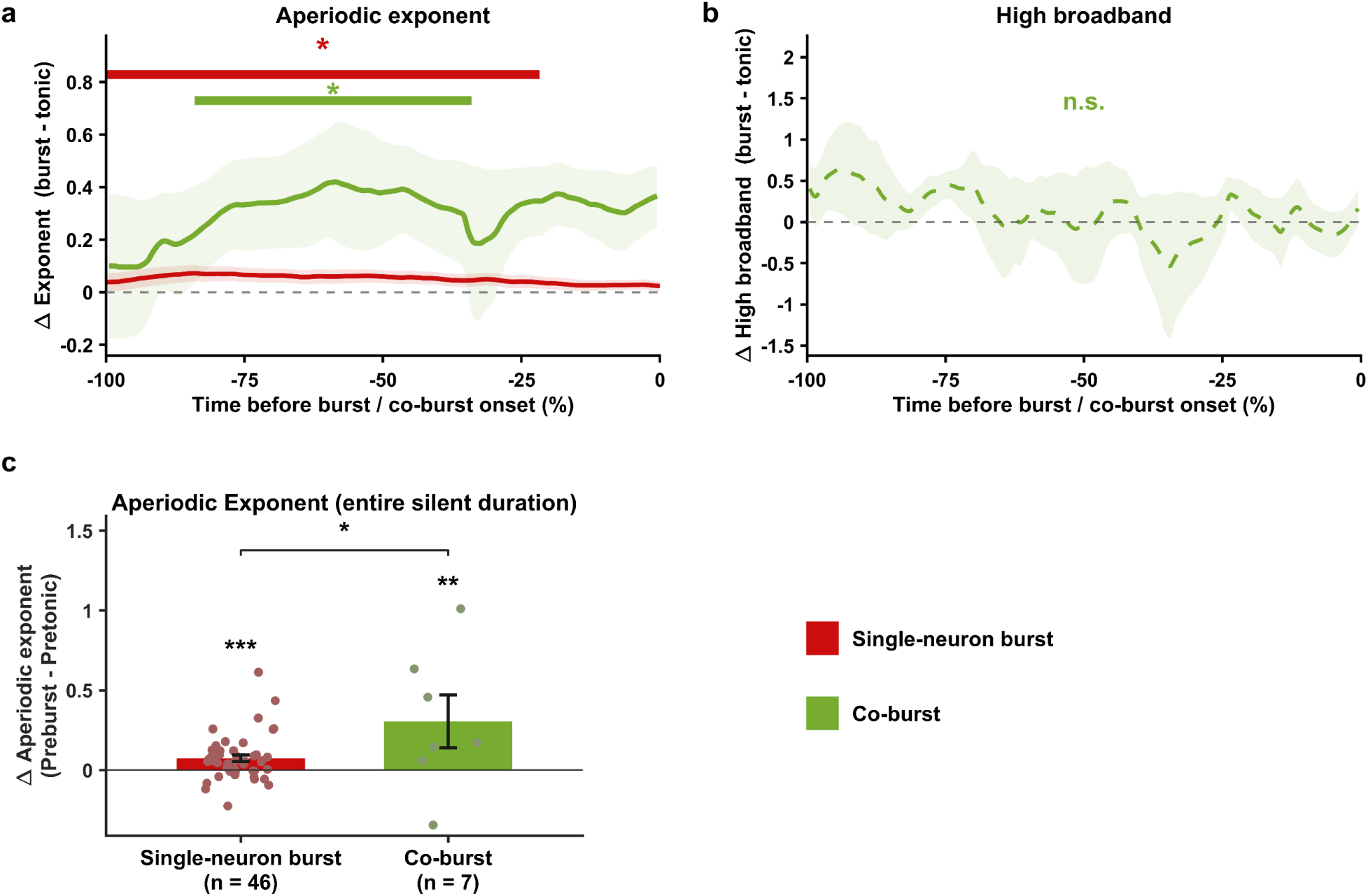
AE increases to a greater magnitude leading up to co-bursting. a. Change in aperiodic exponent (burst - tonic) as a function of time before burst or co-burst onset (-100% to 0) for pre-co-burst vs. matched tonic periods (green; n = 7) and pre-burst vs. pre-tonic periods (red, same as Fig 3a). Horizontal bars denote significant time windows (cluster permutation test). b. Change in high-frequency broadband power for pre-co-burst vs. matched tonic periods (n = 7), which showed no significant difference (n.s.). c. Average change in aperiodic exponent during bursting for single neurons (red) and co-burst populations (green); the increase is larger before co-bursting (*). Dots are individual units or populations; bars show the mean ± error bars.

## DISCUSSION

We characterized putative LTS bursting neurons in the human pulvinar, a higher-order thalamic nucleus, at quiet wakefulness and during task performance. We note that prior human data on thalamic bursting have been primarily from acute intraoperative recordings in the asleep or anesthetized state, with the exception of recordings during a cognitive task (Kim, Ohara and Lenz, 2009).We obtained concurrent LFP recordings and found that the aperiodic exponent was higher in the silent period preceding bursts than in duration-matched silent periods preceding tonic spikes. This effect held after controlling for the firing rate and synchrony of concurrently recorded neurons. The exponent therefore tracked pre-burst E:I dynamics in a manner consistent with intracellular recordings - sustained hyperpolarization. Moreover, greater peak aperiodic exponent and longer silent period durations were associated with larger bursts, consistent with the known graded nature of burst priming, whereby deeper and longer hyperpolarization produces larger bursts (Alitto *et al*., 2019). Findings generalized across behavioral states, with two differences: single-neuron bursting was more frequent at quiet wakefulness, whereas co-bursting was detected at the population level during task performance but not at rest. Recent work in awake monkeys showed that pulvinar bursting is predictive of behavioral performance in an attention task (Boshra *et al*., 2026). Pulvinar relay cells lack local recurrent excitation, so their output is determined largely by afferent excitation and inhibition (Halassa and Sherman, 2019). Inhibition arises from the thalamic reticular nucleus and extrathalamic sources, which is the source of hyperpolarization preceding awake bursts (Barthó *et al*., 2004). Co-bursting is in turn driven by shared excitatory input from cortical populations, consistent with our findings that co-bursting occurred on slow timescales (∼100 ms). In sum, our recordings revealed, to our knowledge, first human single-neuron evidence in support of the E:I hypothesis of the aperiodic exponent, which has otherwise rested on pharmacological and state comparisons (Gao, Peterson and Voytek, 2017; Lendner *et al*., 2020).

## METHODS

### Participants

Intracranial stereo-electroencephalography (SEEG) data were recorded from patients with drug-resistant epilepsy. Thalamic sampling was performed strictly by clinical indication and was prescribed on a case-by-case basis during an epilepsy surgery conference. Single unit recordings were obtained with hybrid electrodes consisting of Behnke-Fried microwires (Ad-Tech Medical, Racine WI) in addition to the standard clinical macro-contacts. As such, research participation did not influence the location or number of SEEG implants. Written informed consent was obtained in all subjects, and our research procedures were approved by our local Institutional Review Board (IRB).

#### Task cohort

Ten subjects (age 44 ± 11.9 years, mean ± s.d.; range 21–64; 8 male) were included. They performed a cognitive task (affective task switching, ATS).

#### Rest cohort

Eight recordings from four subjects (43.3 ± 8.3 years; range 31–49; 4 male).

#### Overall sample

Across both datasets, we studied 11 unique subjects (44.1 ± 11.3 years; range 21–64; 9 male). Three subjects contributed to both the task and rest recordings.

### Behavioral States

#### Task

Subjects viewed naturalistic faces while making binary judgments of either affective features (happy vs. angry) or non-affective features (female vs. male). They first performed single-task blocks and then mixed-task blocks, in which they performed 3-7 consecutive repeat trials of the same task before switching. .

#### At Rest

Patients were awake and at rest but were not involved in any cognitive tasks during this time.

### Data Acquisition and Preprocessing

We used depth electrodes (Ad-Tech) with 6-16 platinum macrocontacts (2.41 mm contact length, 3-5 mm spacing) to record LFP data at 4 kHz (Natus System). We recorded single units from the Behnke-Fried depth (Ad-Tech Medical, Racine WI) electrode at 32kHz. For the recording of micro and macro channels, we used an Atlas (Neuralynx Inc., Bozeman, Montana, USA) acquisition system that allows us to simultaneously record from both the micro and macro channels.

We preprocessed the LFP data by demeaning and detrending, then notch-filtering at 60 Hz (2nd-order IIR filter) and downsampling to 1kHz. We performed bipolar referencing across adjacent contacts within the same electrode. For current analysis, we used only the bipolar “virtual contact” closest to the Behnke-Fried (BF) microwire. For single units analysis, we common-average-referenced all 8 channels of the wire bundle to eliminate noise before using a sorting algorithm to extract single units.

### Spike Sorting and Detection

We performed spike detection and sorting with a modified version of the Osort algorithm(Rutishauser, Schuman and Mamelak, 2006; Buccino *et al*., 2020). In total, we isolated 119 neurons from 10 task sessions and 95 isolated single neurons from the rest of the cohort. We detected bursts from each neuron’s raw spike train.

### Burst detection

A burst was identified as at least 2 or more spikes with a first ISI ≤ 7ms and subsequent intervals ≤ 15 ms preceded by 100ms of silence since the first burst spike(Domich, Oakson and Steriade, 1986; Ramcharan, Gnadt and Sherman, 2000). We applied a per-neuron inclusion criterion requiring at least 10 bursts per neuron over the entire recording. The task lasted roughly an hour, whereas rest lasted around 30 minutes. 83.2% of task-state neurons burst at least once, and 38.7% met the minimum 10-burst criterion; for rest neurons, 72.6% had at least 1 burst, and 28.4% met the minimum burst threshold.

### LTS Criteria

Each neuron with ≥ 10 bursts was further checked for 2 criteria: 1. Successive intervals inside bursts must consistently increase, and 2. The first ISI should decrease with increasing burst size. A neuron satisfying either of these was classified as an LTS unit (Domich, Oakson and Steriade, 1986; Ramcharan *et al*., 2000; Ramcharan, Gnadt and Sherman, 2000).

### Aperiodic exponent

For each bipolar referenced macro channel, the continuous LFP was decomposed into logarithmically spaced frequency bands using a zero-phase FIR band-pass filter. For each band, we applied the Hilbert transform to obtain the analytic signal. We then obtained instantaneous power by squaring the magnitude. To account for increasing bandwidths, we normalized power in each band by its bandwidth. At each time point, log-transformed power was regressed using linear regression restricted to bands with center frequencies between 10-50Hz(Gao, Peterson and Voytek, 2017; Donoghue *et al*., 2020). We used the absolute value of the fitted aperiodic slope as the aperiodic exponent.

### Broadband power

We derived broadband power from the same band-limited analytic signal used for the aperiodic exponent. We converted instantaneous power in each band to a one-sided power spectral density by normalizing by the integral of the filter’s squared two-pass magnitude response. We retained the 13 bands with center frequencies of 70 Hz or higher (centers from 70.7 to 200 Hz), converted the PSD to decibels (10·log₁₀), and averaged across these bands at each time point (Manning *et al*., 2009; Saez *et al*., 2018)

### Aperiodic Exponent before Bursts

For each burst window, the pre-burst window period is the silent interval between the preceding spike and the burst’s first spike. Tonic windows were the equivalent intervals between consecutive non-burst spikes, with the same 100-500 ms limits as the pre-burst silence window limits. Because window length varies for each burst, we normalized the silent period in time (– 100% to 0% of the silent interval) and then averaged within each bin, making different silent durations comparable. We subtracted the mean tonic trace from the burst trace.

For each neuron, we drew 1000 resamples. For each resample, we randomly swapped the tonic and burst labels. We used a cluster-based permutation test (Maris and Oostenveld, 2007) to test the burst-minus-tonic difference averaged across neurons. Bins were clustered if their absolute difference exceeded the top 16% of absolute permuted differences, pooled over permutations and bins. The cluster statistic was the sum of these cluster bins. The null was the largest cluster statistic in each permutation. Clusters outside the 2.5th–97.5th percentiles were significant (two-tailed, p = 0.05).

### Correlations

Since the exponent increases in the pre-burst silence, we tested if the effect is graded. That is, whether longer duration in the pre-silent period and the peak value of the exponent correlate with the burst size. For each LTS burst, we calculated the peak exponent, the pre-burst silence duration (ms), and the number of spikes during the burst. For each neuron that met the minimum 10-burst criterion, we computed Spearman’s correlation between each measure and burst size. Only neurons with variable burst sizes were included in this analysis (task: 39 of 46 neurons; rest: 27 of 27 neurons), as the correlation between silence duration and burst size is undefined when burst size does not vary.

### Mixed Effects Model

To test whether the aperiodic exponent differs between bursts and tonic spikes while accounting for overlap in silent windows, as well as firing rate and population synchrony, we fit a linear within-between mixed-effects model (Laird and Ware, 1982) separately for the task and rest cohorts. The outcome is the mean absolute aperiodic exponent. Fixed effects included neuronal firing state (burst vs tonic), and two control variables, firing rate and pairwise spike synchrony. Random effects included a random intercept and a burst-state slope per neuron. We matched tonic windows to the number of burst windows, with the same silent periods and no overlaps across windows. We also ensured that the pre-tonic period for any neuron was not overlapped with the pre-burst periods of another neuron. We tested the state effect with a likelihood-ratio test comparing the full model to a null model with only the random intercept and fixed effects. Models were fitted with fitlme in MATLAB using maximum likelihood for the likelihood-ratio test.

### Co-bursting

Within a simultaneously recorded population, a conditional-discharge-rate cross-correlogram(Liu *et al*., 2026) was constructed between its burst onsets (the trigger train) and the burst-population train formed by the summed burst onsets of all *other* recorded neurons. We normalized the count of triggered events at each lag by the number of trigger bursts and the bin size, using 1 ms bins over a ±2 s lag window, resulting in a rate expressed in events per second. Positive lags represent triggered events occurring after the trigger onset, with lag 0 corresponding to the same bin.

The observed CCH was compared against a null hypothesis that was constructed by jittering the triggered train(Harrison and Geman, 2009; Amarasingham *et al*., 2012; Liu *et al*., 2026). For each burst onset of every other neuron, we selected an independent integer number of ms from a uniform distribution over ±δ (excluding zero) and shifted the onset by that amount. We used wrap-around so the total number of events was preserved and the trigger train was unaffected. We then reconstructed the CCH in exactly the same way as for the observed data. Seven values of jitter width were tested (δ = 10, 20, 50, 100, 200, 400, 800 ms). We smoothed all observed and surrogate CCHs with a Gaussian kernel (SD = 0.2δ) under edge-replicated padding, and restricted the analysis to |lag| <= 1500ms to avoid edge effects. For each surrogate, the maximum smoothed CCH over the evaluation region defined the null distribution of maxima; we used the one-sided 97.5th percentile threshold. A CCH was significant if any evaluation lag exceeded this threshold, and the empirical p-value was calculated as the fraction of surrogate maxima at least as large as the observed maximum.

### Statistics

We performed statistical analyses using custom scripts in MATLAB (R2024b). All tests were non-parametric except the mixed-effects model, which assumes normally distributed residuals and random effects. We corrected multiple comparisons with the False Discovery Rate (FDR) (Benjamini and Hochberg, 1995), applied to each condition rather than across brain states. We used a threshold of P<0.05, and all reported significant results survived FDR Correction (q<0.05).

## REFERENCES

1. Alitto, H. et al. (2019) “The Augmentation of Retinogeniculate Communication during Thalamic Burst Mode,” The Journal of Neuroscience, 39(29), pp. 5697– 5710. Available at: 10.1523/JNEUROSCI.2320-18.2019.

2. Amarasingham, A. et al. (2012) “Conditional modeling and the jitter method of spike resampling,” Journal of Neurophysiology, 107(2), pp. 517–531. Available at: 10.1152/jn.00633.2011.

3. Barthó, P. et al. (2004) “Characterization of Neocortical Principal Cells and Interneurons by Network Interactions and Extracellular Features,” Journal of Neurophysiology, 92(1), pp. 600–608. Available at: 10.1152/jn.01170.2003.

4. Bates, D. et al. (2015) “Fitting Linear Mixed-Effects Models Using **lme4**,” Journal of Statistical Software, 67(1). Available at: 10.18637/jss.v067.i01.

5. Benjamini, Y. and Hochberg, Y. (1995) “Controlling the False Discovery Rate: A Practical and Powerful Approach to Multiple Testing,” Journal of the Royal Statistical Society Series B: Statistical Methodology, 57(1), pp. 289–300. Available at: 10.1111/j.2517-6161.1995.tb02031.x.

6. Boshra, R. et al. (2026) “Higher-order thalamic bursts are drivers of attention control,” Neuron, p. S0896627326004174. Available at: 10.1016/j.neuron.2026.05.025.

7. Buccino, A.P. et al. (2020) “SpikeInterface, a unified framework for spike sorting,” eLife, 9, p. e61834. Available at: 10.7554/eLife.61834.

8. Buzsáki, G., Anastassiou, C.A. and Koch, C. (2012) “The origin of extracellular fields and currents — EEG, ECoG, LFP and spikes,” Nature Reviews Neuroscience, 13(6), pp. 407–420. Available at: 10.1038/nrn3241.

9. Diehl, G.W. and Redish, A.D. (2024) “Measuring excitation-inhibition balance through spectral components of local field potentials.” Neuroscience. Available at: 10.1101/2024.01.24.577086.

10. Domich, L., Oakson, G. and Steriade, M. (1986) “Thalamic burst patterns in the naturally sleeping cat: a comparison between cortically projecting and reticularis neurones.,” The Journal of Physiology, 379(1), pp. 429–449. Available at: 10.1113/jphysiol.1986.sp016262.

11. Donoghue, T. et al. (2020) “Parameterizing neural power spectra into periodic and aperiodic components,” Nature Neuroscience, 23(12), pp. 1655–1665. Available at: 10.1038/s41593-020-00744-x.

12. Gao, R., Peterson, E.J. and Voytek, B. (2017) “Inferring synaptic excitation/inhibition balance from field potentials,” NeuroImage, 158, pp. 70–78. Available at: 10.1016/j.neuroimage.2017.06.078.

13. Gutierrez, C. et al. (2001) “Dynamics of low-threshold spike activation in relay neurons of the cat lateral geniculate nucleus,” The Journal of Neuroscience: The Official Journal of the Society for Neuroscience, 21(3), pp. 1022–1032. Available at: 10.1523/JNEUROSCI.21-03-01022.2001.

14. Halassa, M.M. and Sherman, S.M. (2019) “Thalamocortical Circuit Motifs: A General Framework,” Neuron, 103(5), pp. 762–770. Available at: 10.1016/j.neuron.2019.06.005.

15. Harrison, M.T. and Geman, S. (2009) “A Rate and History-Preserving Resampling Algorithm for Neural Spike Trains,” Neural Computation, 21(5), pp. 1244–1258. Available at: 10.1162/neco.2008.03-08-730.

16. Jahnsen, H. and Llinás, R. (1984) “Electrophysiological properties of guinea-pig thalamic neurones: an in vitro study.,” The Journal of Physiology, 349(1), pp. 205–226. Available at: 10.1113/jphysiol.1984.sp015153.

17. Kim, J.H., Ohara, S. and Lenz, F.A. (2009) “Mental Arithmetic Leads to Multiple Discrete Changes From Baseline in the Firing Patterns of Human Thalamic Neurons,” Journal of Neurophysiology, 101(4), pp. 2107–2119. Available at: 10.1152/jn.91087.2008.

18. Laird, N.M. and Ware, J.H. (1982) “Random-effects models for longitudinal data,” Biometrics, 38(4), pp. 963–974.

19. Lendner, J.D. et al. (2020) “An electrophysiological marker of arousal level in humans,” eLife, 9, p. e55092. Available at: 10.7554/eLife.55092.

20. Liu, X. et al. (2026) “Entrained cortical delta–spindle activity, not periodic synchrony, prevents arousal by NREM thalamic bursts,” Communications Biology, 9(1), p. 285. Available at: 10.1038/s42003-026-09565-3.

21. Lu, S.M., Guido, W. and Sherman, S.M. (1992) “Effects of membrane voltage on receptive field properties of lateral geniculate neurons in the cat: contributions of the low-threshold Ca2+ conductance,” Journal of Neurophysiology, 68(6), pp. 2185–2198. Available at: 10.1152/jn.1992.68.6.2185.

22. Manning, J.R. et al. (2009) “Broadband Shifts in Local Field Potential Power Spectra Are Correlated with Single-Neuron Spiking in Humans,” The Journal of Neuroscience, 29(43), pp. 13613–13620. Available at: 10.1523/JNEUROSCI.2041-09.2009.

23. Maris, E. and Oostenveld, R. (2007) “Nonparametric statistical testing of EEG- and MEG-data,” Journal of Neuroscience Methods, 164(1), pp. 177–190. Available at: 10.1016/j.jneumeth.2007.03.024.

24. Miller, K.J., et al. (2009) “Power-Law Scaling in the Brain Surface Electric Potential,” PLoS Computational Biology. Edited by O. Sporns, 5(12), p. e1000609. Available at: 10.1371/journal.pcbi.1000609.

25. Ramcharan, E.J. et al. (2000) “Cellular Mechanisms Underlying Activity Patterns in the Monkey Thalamus During Visual Behavior,” Journal of Neurophysiology, 84(4), pp. 1982–1987. Available at: 10.1152/jn.2000.84.4.1982.

26. Ramcharan, E.J., Gnadt, J.W. and Sherman, S.M. (2000) “Burst and tonic firing in thalamic cells of unanesthetized, behaving monkeys,” Visual Neuroscience, 17(1), pp. 55–62. Available at: 10.1017/S0952523800171056.

27. Ramcharan, E.J., Gnadt, J.W. and Sherman, S.M. (2005) “Higher-order thalamic relays burst more than first-order relays,” Proceedings of the National Academy of Sciences, 102(34), pp. 12236–12241. Available at: 10.1073/pnas.0502843102.

28. Rutishauser, U., Schuman, E.M. and Mamelak, A.N. (2006) “Online detection and sorting of extracellularly recorded action potentials in human medial temporal lobe recordings, in vivo,” Journal of Neuroscience Methods, 154(1–2), pp. 204–224. Available at: 10.1016/j.jneumeth.2005.12.033.

29. Saez, I. et al. (2018) “Encoding of Multiple Reward-Related Computations in Transient and Sustained High-Frequency Activity in Human OFC,” Current Biology, 28(18), pp. 2889–2899.e3. Available at: 10.1016/j.cub.2018.07.045.

30. Sherman, S. Murray and Guillery, R.W. (2002) “The role of the thalamus in the flow of information to the cortex,” Philosophical Transactions of the Royal Society of London. Series B: Biological Sciences. Edited by P. Adams et al., 357(1428), pp. 1695–1708. Available at: 10.1098/rstb.2002.1161.

31. Sherman, S.M. (2001) “Tonic and burst firing: dual modes of thalamocortical relay,” Trends in Neurosciences, 24(2), pp. 122–126. Available at: 10.1016/S0166-2236(00)01714-8.

32. Wiest, C. et al. (2023) “The aperiodic exponent of subthalamic field potentials reflects excitation/inhibition balance in Parkinsonism,” eLife, 12, p. e82467. Available at: 10.7554/eLife.82467.

